# Left hemispheric deficit in the sustained neuromagnetic response to periodic click trains in children with ASD

**DOI:** 10.1101/2020.07.24.219410

**Authors:** T.A. Stroganova, K.S Komarov, D.E. Goiaeva, T.S. Obukhova, T.M. Ovsiannikova, A.O. Prokofyev, E.V. Orekhova

## Abstract

Deficits in perception and production of vocal pitch are often observed in people with autism spectrum disorders (ASD), but the neural basis of these abnormalities is unknown. In magnetoencephalogram (MEG), spectrally complex periodic sounds trigger two continuous neural responses – the Auditory Steady State Response (ASSR) and the Sustained Field (SF). It has been shown that the SF in neuro-typical individuals is associated with low-level analysis of pitch in the ‘pitch processing center’ of the Heschl’s gyrus. Therefore, this auditory response may reflect vocal pitch processing abnormalities in ASD. The SF, however, has never been studied in people with these disorders. We used MEG and individual brain models to investigate the ASSR and SF evoked by monaural 40 Hz click trains in 7-13-year-old boys with ASD (N=35) and neuro-typical (NT) boys (N=35). In agreement with the previous research in adults, the cortical sources of the SF in children were located in the left and the right Heschl’s gyri, anterolateral to those of the ASSR. In both groups, the SF and ASSR dominated in the right hemisphere and were higher contralaterally to the stimulated ear. The ASSR increased with age in both NT and ASD children and did not differ between the groups. The SF was moderately decreased in both hemispheres and was markedly delayed and displaced in the left hemisphere in boys with ASD. The SF delay in participants with ASD was present irrespective of their intelligence level and severity of autism symptoms. We suggest that the selective left-hemispheric SF abnormalities in children with ASD reflect a low-level deficiency in pitch processing that may contribute to their difficulties with perception and production of linguistic prosody.

## Background

Autism spectrum disorder (ASD) is a neurodevelopmental disorder, characterized by deficient verbal and non-verbal communication skills, restricted interests, repetitive behaviors, and is often associated with low intelligence [1]. Although atypical development of spoken language is among the core symptoms of ASD, language impairments are highly variable across individuals with ASD, ranging from complete lack of spoken language to incomplete understanding of language pragmatics [2]. Neural origin of spoken language deficiency in people with ASD remains poorly understood.

Neuroimaging studies have shown that atypical development in children with ASD may originate from an early aberration of the temporal cortex maturation, which is specific to the left hemisphere [3]. It remains unknown, however, when and where in the brain a left-hemispheric speech processing deficit arises in individuals with ASD. One of the most intriguing aspect of this problem is whether such a lateralized deficit exists already at the low level of cortical hierarchy – in the regions of the auditory core confined by the Heschl’s gyrus [4] that extracts the periodicity/pitch from the acoustic input. This ‘low-level’ neurofunctional deficit may manifest itself in different ways. Firstly, it might result in atypical frequency encoding of the auditory input that relies on tonotopic neuronal representations of distinct frequency channels, inherited by the primary auditory cortex (A1) from subcortical auditory pathways [5]. Secondly, it may cause abnormal processing of the temporal regularities in the amplitude envelope of a spectrally complex auditory signal. While slow temporal modulations at frequencies from 4-to 16-Hz play an important role in comprehension of speech [6, 7], those of higher frequencies (> ~30 Hz) are essential for perception of vocal pitch [8]. Unlike pure tones, periodic temporal modulation of the complex sound (e.g. repetitive mixture of harmonics or repetitive transients/noise) is not solely place-coded in the A1, but is also processed in another region of the core auditory cortex [9]. Neuronal findings in monkeys [10, 11] and functional neuroimaging in humans [12] have shown that the so-called ‘pitch processing center’ in the auditory cortical hierarchy is localized downstream to A1 in the more anterolateral portion of the Heschl’s gyrus, at the border of the core and the belt auditory cortical areas. Given that the individuals with ASD, regardless of their IQ and general language skills, have abnormalities in perception of pitch in speech signals, as well as difficulties with production of adequately intonated speech [13, 14], a putative low-level neural deficit in encoding of temporal regularity in this clinical group merits careful investigation.

While many studies associate pitch processing primarily with the right auditory cortex (e.g. [15]), there is strong evidence that temporal regularities perceived as a pitch are processed bilaterally, but differently in the left and right hemispheres [16, 17]. The low-level processing of temporal regularities in spectrally complex sound can be investigated by measuring lateralized neural auditory responses to sequences of periodic clicks presented monaurally to the left and right ears. According to findings in healthy adults, the prolonged periodic click trains evoke two types of magnetoencephalographic (MEG) responses in the auditory cortex – the oscillatory auditory steady-state response (ASSR) at the frequency of stimulation and the sustained deflection of the magnetic field (‘Sustained Field’ - SF) [18–21].

Although both the SF and ASSR were previously associated with processing of periodicity/temporal regularity [19, 21], only the 40 Hz ASSR has been examined in individuals with ASD [22–25]. The ASSR is an oscillatory response phase-locked to the onset of each single auditory event, i.e. each click in the train. The maximal ASSR is observed at the stimulation frequency of 40 Hz that is thought to represent the resonance frequency of the A1 neural circuitry [26]. The interest to the 40 Hz ASSR in people with ASD was driven by it’s putative link to local connectivity disturbances in these disorders [25] and numerous reports on its robust reduction in patients with schizophrenia (for review see [27]).

In contrast to schizophrenia research, only a few studies examined the 40 Hz ASSR in ASD, and the results of these studies are inconsistent. Reduction of the ASSR was found either bilaterally in adolescents and adults with ASD [24], as well as in parents of children with ASD [23], or unilaterally in the left hemisphere, both in children with ASD and their fist-degree relatives ([25] and [28] respectively). In addition to uncertainty regarding lateralization of the ASSR abnormality, a recent study by Edgar et al, which included a large sample of participants, found the ASSR to be fairly normal in both hemispheres of children with ASD [29]. Edgar and colleagues also investigated the developmental trajectory of the ASSR and suggested that the discrepancy with the previous results was due to relatively late maturation of this response in humans. They observed that the ASSR was typically weak and unreliable before puberty, which might preclude detection of its potential abnormalities in children with ASD. Another reason for conflicting findings may be the type of stimulation (monaural vs binaural) used in different studies. Monaural as compared to binaural stimulation is known to produce larger hemispheric asymmetry [30] and, therefore, is more suitable to reveal putative hemisphere-specific ASSR abnormality in people with ASD.

Although the ASSR reflects processing of stimulus periodicity within a certain range of frequencies, it is hardly relevant for pitch processing in the auditory cortex. Indeed, the ASSR originates from A1 and gradually decreases and almost disappears at frequencies of 75-150 Hz [31] - the fundamental frequencies of the human voice that mainly determine what is perceived as the pitch of the voice [32]. The higher-order processing associated with perception of pitch is rather reflected in another component – the SF generated in the anterolateral region of Heschl’s gyrus [19, 20, 33]. The SF is a slowly developing DC shift with negative polarity (in the EEG) that is locked to the onset of temporally modulated auditory stimulation and continues throughout its duration [34]. Apart from click trains, the SF can be triggered by any periodic spectrally complex sounds, including amplitude-modulated tones and speech sounds, and is very sensitive to periodicity, which is perceived as a sustained pitch [18, 33]. The ‘pitch processing center’ that generates the SF does not belong to the primary auditory cortex, despite the anatomical proximity and cytoarchitectonic similarity of these two cortical regions [4]. Based on these findings, it has been suggested that the SF reflects the integration of pitch information across the frequency channels of the primary auditory cortex, where the sound frequencies are represented as a tonotopic frequency map [19].

Given the relevance of the SF to pitch processing, it is surprising that this neural response escaped the attention of researchers who studied the 40-Hz ASSR in ASD. Moreover, in the neuro-typical (NT) children, the SF response to the 40 Hz clicks has been described only in one study (mean age of participants was 12 years) but its characteristics (localization, hemispheric asymmetry) have not been analyzed [35]. Therefore, it is not known whether the properties of the SF responses to the 40-Hz clicks in children and adults are similar. Taking into account the protracted maturation of the auditory evoked responses [36–39], the properties of the SF might differ in children and adults. If child and adult SF are homologous, studying this response in children with ASD may reveal the suspected abnormalities in processing of pitch in the ‘pitch processing center’. Moreover, exploration of the SF hemispheric asymmetry using a monaural presentation of periodic clicks may shed light on the hemisphere-specific deficit in pitch processing in ASD.

In this study we investigated both types of auditory evoked fields (AEF) associated with processing of sound periodicity in children with ASD: the ASSR and SF. For a more reliable analysis of hemispheric asymmetries, we presented 40 Hz click trains monaurally to the left and right ears. We recorded auditory responses using MEG and applied individual head models for source analysis. This made it possible to localize the cortical sources of both components and determine the direction of the source current, which is important for the neurophysiological interpretation of the results. Given the lack of knowledge about the properties of the SF in children, we ***firstly*** compared the spatial and temporal characteristics of this auditory response in neuro-typical children and adults. ***Secondly***, we compared the ASSR and SF in the NT children and those with ASD. We expected that if there is a difference between the groups, it might be specific to a particular type of response and/or confined to one hemisphere. ***Thirdly***, we examined whether inter-individual variations of the ASSR and SF – the components associated with low-level processing of non-speech periodic sound - correlate with cognitive deficit and the core symptoms of autism in children with ASD.

## Methods

### Participants

The study included 35 boys with ASD aged 7-13 years and 35 NT boys of the same age range. The participants with ASD were recruited from rehabilitation centers affiliated with the Moscow University of Psychology and Education, and were assessed by the same experienced licensed psychiatrist. None of the NT participants had known neurological or psychiatric disorders. All participants had normal hearing according to medical records. In children, IQ has been evaluated through standard scores on K-ABC subscales (Simultaneous and Sequential), as well as by calculating Mental Processing Index (MPI) [40]. Presence and severity of autism symptoms has been evaluated using Russian translations of three parental questionnaires: Social Responsiveness Scale for children [41], Autism spectrum Quotient (AQ) for children [42], and Social Communication Questionnaire (SCQ-Lifetime) [43]. Table 1 presents characteristics of the pediatric samples. To compare child and adult auditory responses to click trains we included in the study 10 NT adults (ages 22-36 years, mean (SD) =28.5 (4.1), 5 females).

**Table 1.** Characteristics of the pediatric samples (mean, SD, range, group differences).

|  | ASD | NT | p | T |
| --- | --- | --- | --- | --- |
| Age | 9.69±1.5 | 10.08±1.5 | 0.29 | 1.09 |
| N<sub>NT/ASD</sub> =35/35 | (7.2-12.3) | (7.3-13.0) |  |  |
| Sequential IQ | 78.9±19.1 | 106.8±13.7 | <1e-6 | 6.93 |
| N<sub>NT/ASD</sub> =35/32 | (49-127) | (77-134) |  |  |
| Simultaneous IQ | 89.2±22.0 | 123.3±15.3 | <1e-6 | 7.43 |
| N<sub>NT/ASD</sub> =35/32 | (43-134) | (91-151) |  |  |
| MPI Standard IQ | 84.3±22.0 | 121.2±12.6 | <1e-6 | 8.51 |
| N<sub>NT/ASD</sub> =35/32 | (45-128) | (94-145) |  |  |
| SRS | 104.8±23.9 | 45.2±22.6 | <1e-6 | -10,4 |
| N<sub>NT/ASD</sub> =34/32 | (57-157) | (3-91) |  |  |
| SCQ-life | 24.7±5.4 | 6.6±4.2 | <1e-6 | -15,0 |
| N<sub>NT/ASD</sub> =33/32 | (11-34) | (0-19) |  |  |
| AQ | 90.3±12.2 | 53.7±13.0 | <1e-6 | -11,7 |
| N<sub>NT/ASD</sub> =33/32 | (61-118) | (23-76) |  |  |

‘N<sub>NT/ASD</sub>’ indicates the number of participants of the corresponding groups for whom the data were available.

The investigation was approved by the Ethical Committee of the Moscow University of Psychology and Education. All participants and/or their caregivers provided their verbal assent to participate in the study and were informed about their right to withdraw from the study at any time during testing. The adult participants and guardians of all children gave written informed consent after the experimental procedures had been fully explained.

### Auditory stimuli

The stimuli - 500 ms trains of 40 Hz clicks - were delivered via plastic ear tubes inserted in the ear channels. The intensity level was set at 60 dB SPL. The duration of each click was 2ms and the stimulus onset asynchrony was 25 ms, resulting in a train of 20 clicks. Intervals between the trains were fixes at 1000 ms. The stimuli were organized in two blocks, each containing 100 click trains presented to one ear. The order of the ‘left’ and ‘right’ blocks was counterbalanced between subjects. The experiment lasted for approximately 5 minutes. Participants were instructed to ignore the auditory stimulation and watched a silent video of their choice during the experiment.

Waveform and spectral composition of the stimulus are presented in Figure 1. The signal energy is concentrated at 40 Hz (*f_0_*, fundamental frequency), as well as at the higher frequency harmonics of comparable physical intensity. Periodicity (‘pitch’) in such spectrally complex sound can be analyzed by the auditory system based either on the lowest harmonic present - *f_0_*, or on the highest common devisor of the sound’s harmonics (*f_sp_*) [16, 44].

**Figure 1.**
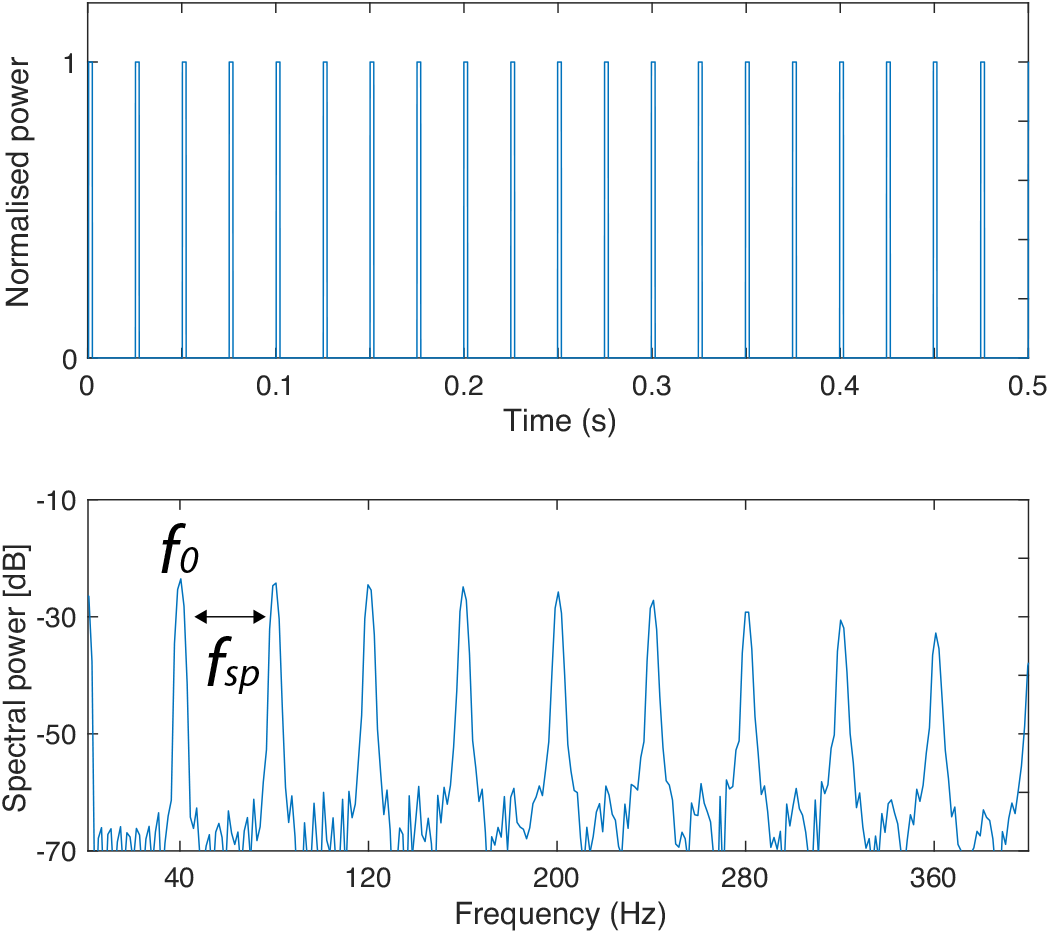
Waveform (upper panel) and power spectrum (lower panel) of the 40 Hz click train. Periodicity (‘pitch’) can be analyzed by the auditory system based either on the fundamental frequency (*f*_0_), or on the common devisor of the sound’s harmonics (*f_sp_*).

### MEG acquisition and preprocessing

MEG was recorded in a sitting position in a neuromagnetically-shielded room using a 306-channel MEG scanner (Vectorview, Elekta-Neuromag) that comprises 204 orthogonal planar gradiometers and 102 magnetometers in 102 locations above the participant’s head. An electrooculogram (EOG) was recorded using four electrodes placed at the outer canthi of the eyes as well as above and below the left eye. To monitor the heartbeats one ECG electrode was placed at the manubrium sterni and the other one at the mid-axillary line (V6 ECG lead). The signal was sampled at 1000 Hz and filtered on-line with a bandpass filter of 0.1–330 Hz.

Prior to the MEG recording, the positions of HPI coils, fiducial points and additional points on the head and face were digitized using the 3D digitizer ‘FASTRAK’ (Polhemus, Colchester, VT, US). The subject’s head position inside the MEG helmet was assessed every 4 ms. The temporal signal space separation (tSSS) and movement compensation options implemented in MaxFilter software (Elekta-Neuromag) were used to suppress environmental noises and compensate for head movements. The data were converted to standard head position (x = 0 mm; y = 0 mm; z= 45 mm).

After the signal-space separation, magnetometers and gradiometers contain nearly equivalent information [45]. We used only gradiometers for analysis. The MEG data preprocessing was performed using MNE-python software [46] and customary scripts. The raw data were notch-filtered at 50, 100 and 150 Hz. To detect the sustained component of the auditory response we have chosen not to apply an off-line high-pass filter. Independent component analysis performed on the continuous data was used to reject components associated with eye movements, heartbeats and muscle artifacts. The data segments characterized by too low (<1e-13 T/m) or too high (>4000e-13 T/m) amplitudes were excluded from ICA and the following analysis. Number of rejected components per subject did not differ in the NT and ASD groups (NT: mean=3.8, STD=0.96; ASD: mean=3.9, STD=1.22’ p=0.7). The cleaned signal was epoched from −500 to +1000 ms relative to the click train onset. We then rejected the epochs contaminated by occasional SQUID ‘jumps’, based on thresholding the amplitude of the 100 Hz high-passed signal at 3 standard deviations from the mean (the approach used in Fieldtrip; [47]). The trials were then visually inspected to remove remaining epochs contaminated by bursts of muscle activity. The minimal number of the clean epochs per subject and condition was 57 (range 57-97). The mean number of epochs was 89/89 (left/right ear stimulation) in adults, 86/88 in NT children and 77/77 in children with ASD. We visually inspected averaged event-related fields to identify subjects who were lacking visible brain responses to sound. All subjects, including children with ASD, demonstrated sizable responses to the stimulation onset.

### Source localization

To obtain individual source models we used T1-weighted MRIs acquired on a 1.5 T Toshiba ExcelArt Vantage scanner (TR =12 ms, TE = 5 ms, flip angle = 20°, slice thickness = 1.0 mm, voxel size = 1.0 × 1.0 × 1.0 mm3). Cortical reconstruction and volumetric segmentation were performed with the Freesurfer image analysis suite [48]. The gray-matter segment was used to construct a continuous triangular high-density mesh representing the neocortex. The following analysis was performed using the Brainstorm software (Tadel et al., 2011). The MEG-MRI co-registration was performed using six reference points (nasion, left and right preauricular points, anterior commissure, posterior commissure and interhemispheric cleft) and the additional points. For every subject the Freesurfer cortical meshes were downsampled to have 15,000 vertices. To compute brain models we implemented Brainstorm method ‘Overlapping spheres’ which fits one local sphere for each sensor [49]. Source reconstruction was performed using the standardized low-resolution brain electromagnetic tomography (sLORETA) [50]. Noise covariance was estimated in −500 ms to 0 ms time interval preceding the stimulation onset. To facilitate comparison between subjects, the individual sLORETA results were morphed to the ‘Collins 27’ template brain provided by Brainstorm.

### Anatomical labels

Sources of the 40Hz ASSR and SF were estimated within the left and the right auditory cortex. Based on the previous studies that used periodic auditory stimuli [19, 20, 51] we have limited the search for cortical sources by the anatomical labels that overlapped or adjoined the left and right auditory cortices: planum temporale of the superior temporal gyrus, transverse temporal sulcus, transverse temporal gyrus of Heschl, planum polare of the superior temporal gyrus, inferior segment of the circular sulcus of the insula, lateral superior temporal gyrus and posterior ramus of the lateral sulcus [52].

### 40 Hz ASSR

For the ASSR analyses the inverse operator was applied to single epochs. We then calculated 40 Hz inter-trial phase coherence (ITPC) --- a measure of phase consistency over trials, with ITPC = 1 reflecting maximal phase consistency across trials and ITPC = 0 - maximal phase variability across trials. It has previously been shown that ITPC is a more reliable measure compared to the total power in the same frequency range [53], that is also more sensitive to developmental changes in the 40 Hz ASSR [29]. To calculate ITPC we performed time-frequency analysis using multitapers with 200 ms sliding time window moving with 0.01 s steps. The frequency smoothing parameter (‘tapsmofrq’) was set to 4. For each subject and stimulation condition ITPC values were averaged in frequency range of 38 – 42 Hz and in the interval between 180 to 500 ms after the click train onset (further called ‘40 Hz ITPC’). To estimate the NT vs ASD differences, for each child we calculated the 40 Hz ITPC averaged over 10 vertices that had the highest ITPC values in the combined group of children. To estimate random 40 Hz ITPC values we computed the average baseline ITPC in −500 to 0 ms interval in the same locations.

To determine cortical localization of the ASSR, for each participant we estimated MNI coordinates of the vertices with the maximal 40 Hz ITPC values in the left and right hemispheres, contralateral to the stimulated ear. For this analysis we excluded subjects with low SNR, who had the 40 Hz ITPC values lower than 0.18 – the maximal baseline value observed across subjects and conditions. Individual coordinates were then used to explore the differences in cortical localizations of the ASSR between the ASD and NT groups, as well as between the ASSR and SF in each experimental group.

### Sustained Field

The signal was first averaged over epochs and the individual timecourses were obtained for each vertex source and baseline-corrected using −500 to 0 ms pre-stimulus baseline. To find coordinates of the left and right SF maxima, we calculated average absolute current amplitude in the 200-500 ms interval, where the SF was present in both children and adults. The groups average SF coordinates (NT adults, NT children, ASD children) were calculated by averaging the individual SF coordinates across participants.

To define the SF spatial region for the group analysis of the SF amplitude, we averaged absolute amplitudes in the 200-500 ms interval over all children (NT + ASD) or adults and found 30 vertices with the greatest resulting values. The individual SF timecourses were then calculated as an average across these 30 vertices. Since the shift in polarity indexes a distinct neural response, to calculate the SF timecourses we retained sign of the activation [54]. Visual inspection of the results demonstrated that the direction of the SF current in the auditory cortex had predominantly negative polarity. Being similar in the waveform, in some vertices the SF could have an opposite - positive – polarity. The polarity changes were observed in different degree in different participants. These individual differences might reflect anatomical features of individual cortices. Indeed, there is a considerable inter-individual variability in morphology of the auditory cortex: approximately a third of the individuals have two Heschl’s gyri [55, 56. To avoid signal cancellation, the averaging took into account the polarity mismatches {Mamashli, 2017 #124]. This was done by flipping polarity of the timecourses in those vertices that were anti-correlated with the subject’s grand average in the 30 vertices in the 0-500 ms time interval. Since the SF source current had a predominantly negative sign, in all cases this resulted in flipping of occasional positive-polarity timecourses before final averaging. The number of the flipped single-vertex timecourses did not differ in children with ASD and NT children (left hemisphere: t=-0.05, p=0.95; right hemisphere: t=0.26, p=0.79).

For each subject, the averaged signal was low-passed at 9 Hz to filter out 40 Hz ASSR and other high-frequency activity. For the time course analysis we then calculated averaged SF amplitude in 4 consecutive intervals: 150-250 ms, 250-350 ms, 350-450 ms and 450-550 ms.

### Statistical analysis

Statistical analysis was performed in STATISTICA (*TIBCO Software Inc*). Repeated measures ANOVA was used to test for the effects of Contra/Ipsi laterality of the stimulation, Hemisphere and Time (for the SF) and their interactions with Group. Greenhouse-Geisser correction was applied when appropriate. We calculated Pearson correlations for approximately normally distributed values (SF source amplitudes) and Spearman correlations when the distributions deviated from normal in at least one of the experimental groups (40 Hz ITPC).

## Results

### Psychometric results

#### IQ

Children with ASD had significantly lower IQ than NT children (Table 1.). Variability of MPI scores was high in the ASD sample and ranged from ‘very low’ to ‘higher than the average’ scores on all three scales.

#### Autism Scores

According to each of the three parents’ questionnaires used, the severity of autism symptoms was significantly higher in the ASD than in the NT sample (Table 1). There were high correlations between all three ‘autism severity’ scales in both the ASD (Pearson correlation coefficients for SRS/SCQ-life: 0.53, SRS/AQ: 0.52, SCQ-life/AQ: 0.58) and the NT (Pearson correlation coefficients for SRS/SCQ-life: 0.60, SRS/AQ: 0.84, SCQ-life/AQ: 0.51) participants. In order to construct a unified ‘Autism Score’ for neuro-behavioral correlation analysis, we reduced dimensionality of the data by extracting the common variance shared by the three questionnaires. Only data from children with ASD was used for constructing the ‘Autism Score’. The values obtained from the three autism scales were z-transformed before performing the PCA. In the subjects who missed the data on some of the scales (N=4), the missing values were substituted by the mean of the available scales. The 1^st^ principle component, which accounted for 71% of the variance in the ASD group, was then used as the integrative ‘Autism Score’. Children with positive Autism Scores scored higher than the average for the ASD group on autism severity, while negative Autism Scores indicated relatively milder autism symptoms. There was a negative correlation between the ‘Autism Score’ and IQ (MPI Standard) (Pearson correlation coefficient =-0.36, p= 0.049).

### The auditory response waveforms in children and adults

To investigate if the SF is present in children of 7-12 years of age, and if it is homological to the ‘adult’ SF, we compared the whole waveform of the auditory evoked response to click trains in the NT children and adults, both in the ‘sensor space’ and in the ‘source space’. To visualize all the main components of the auditory response – the transient components, the ASSR and SF - the low-pass filter was set at 100Hz.

Figure 2 displays the grand average sensor-space plots (A, B) and sLoreta timecourses in the SF group maxima within the left and right Heschl’s gyri (C; see Methods for details). There were marked differences in the auditory responses to click trains between the NT children and adults.

**Figure 2.**
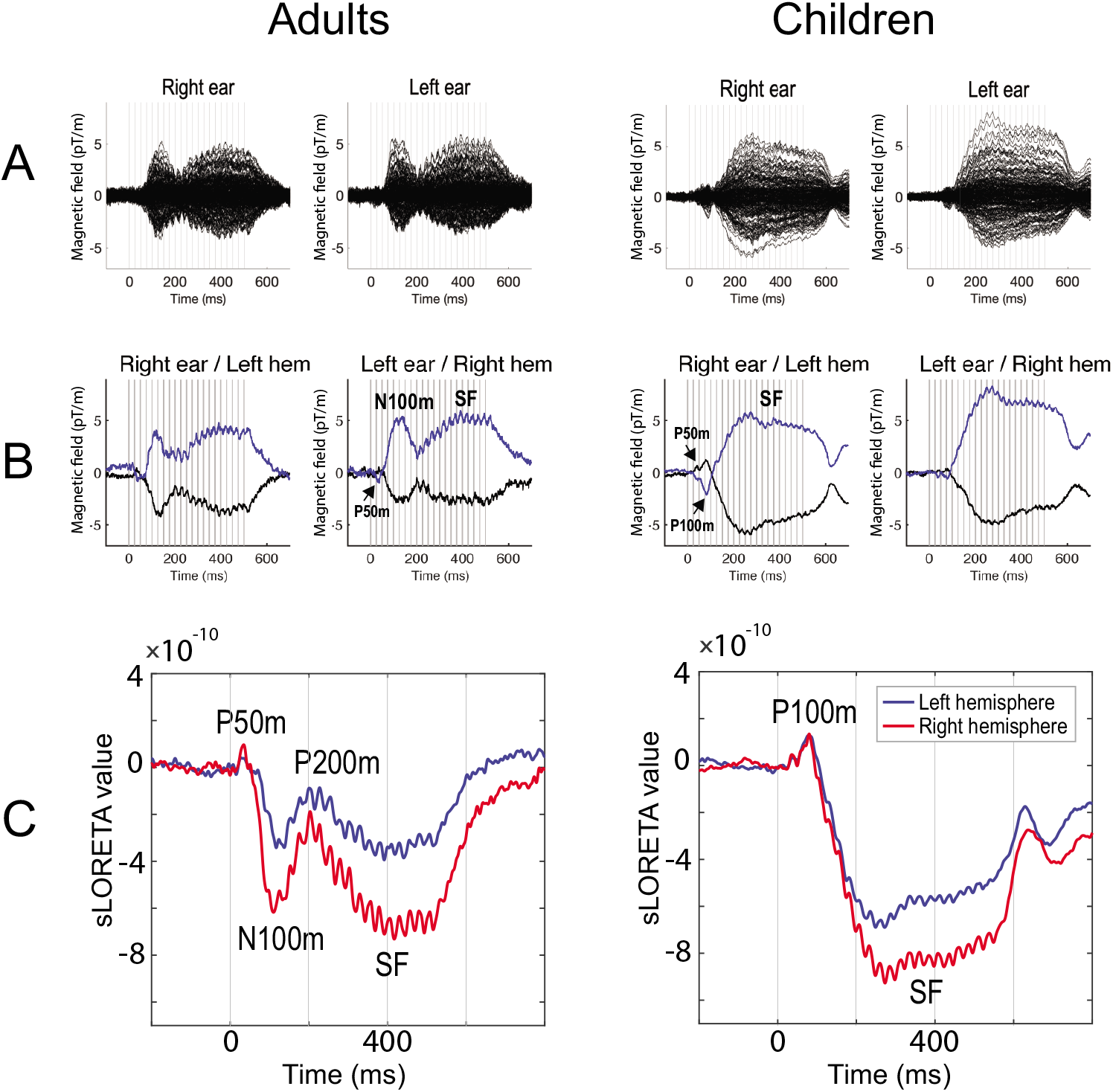
Transient and sustained auditory responses to 40 Hz click trains in NT adults and children: grand average plots. Here and hereafter, zero on the horizontal axes corresponds to the click train onset. The ASSR is present in all plots as an oscillatory 40 Hz steady-state response, which overlaps with the transient components and SF. **A**. ‘Butterfly’ plot of 204 gradiometers for the right- and left-ear stimulation conditions. Vertical gray lines mark onsets of each click in the train. **B.** Auditory responses in the selected left (for the right ear stimulation) and right (for the left ear stimulation) gradiometers. The selected gradiometers are the same in adults and children; they represent the channels measuring maxima of the outgoing/‘positive’ (blue) and incoming/‘negative’ (black) magnetic field flux spatial derivatives. The signal deflections corresponding to the P50m and P100m (in children only) are marked by arrows. **C.** Averaged time courses of the source current at the SF group maxima. Note that direction of the SF source current is negative in both age groups despite marked differences in the successive transient evoked components.

In adults, the stimulation onset evoked a sequence of transient obligatory MEG responses - a small but distinguishable P50m at around 40 ms, followed by a much more prominent N100m of the opposite polarity peaking at around 115 ms after the stimulation onset. The strength of negative current decreased at 200 ms, due to evolving ‘positive’ P200m. These transient components partially overlapped with a slowly developing magnetic field shift (SF), which had the same polarity as the N100m component, reaching its maximal strength at around 400 ms and lasting until the end of stimulation.

In children, the tiny P50m component at about 40 ms was followed by a second deflection of the same positive polarity at 80-84 ms (Figure 2 B, C), which corresponds to the child P100m and is distinct from the P50m and N100m components in adults (Orekhova et al, 2013). The absence of the N100m and the P200m peaks in auditory responses to clicks and tones is typical for children before adolescence [36, 37, 39]. Despite the striking developmental difference in morphology of the transient components, the SF in children resembled that in adults: in both age groups its sources in the auditory cortex had the same direction of current and comparable magnitude (Figure 2 C). Moreover, in response to contralateral stimulation, the SF dominated in the right hemispheres in both age groups. Visibly faster development of the SF in children compared to adults might be explained by the lack of the adult P200m component that masks the early segment of the SF response. In both children and adults, the 40 Hz ASSR overlapping with the is clearly visible on sensors (Figure 2 A, B) and in the source space (Figure 2 C).

### Source localization of the 40Hz ASSR and SF in the auditory cortex

To compare cortical localizations of the ASSR and SF, for each subject we calculated the MNI coordinates of the sources with the maximal 40 Hz ITPC in the 180-500 ms range and those with the maximal integrated SF amplitude in the 200-500 ms range. Only cortical responses that were contralateral to the stimulated ear were used for this analysis. Since localization results are highly sensitive to SNR, for ASSR localization analysis we included only subjects with the 40 Hz ITPC values higher than 0.18 (the maximal baseline value across subjects and conditions): 27 NT and 24 ASD children in case of the right ear stimulation, and 32 NT and 30 ASD children for the left ear stimulation. For the SF localization analysis we included subjects who had visually detectable SW responses in the hemisphere contralateral to the stimulated ear (35 NT and 35 ASD children for the left ear stimulation, 33 NT and 29 ASD children for the right ear stimulation). Both sets of data that fulfilled the described criteria were available for all adults for both hemispheres, for 26 NT and 21 ASD children for the left hemisphere, and for 32 NT and 28 ASD children for the right hemisphere.

The MNI coordinates of the SF and the 40 Hz ITPC maxima in the three groups of participants are shown in Table 3 and visualized in Figure 3. In the NT children, in both hemispheres, the SF source was located anterior, lateral and inferior to that of the ASSR (paired T-tests, all p’s<0.05). The same relative positions of the ASSR and SF sources in adult participants were previously described by Keceli and colleagues [20] using single dipole modeling. For comparison purposes, Table 3 gives original Talairach and estimated MNI coordinates of the SF and ASSR, reported by Keceli et al [20]. In our adult sample, the SF maxima were located anterior and inferior to those of the ASSR in both hemispheres (paired T-test, all p’s<0.05), while the lateral shift along the X axis was not significant, possibly because of the small sample size. In children with ASD, the differences in cortical localization between the SF and ASSR sources were in the same direction as in the NT children, though not always significant (Right X coordinate: T=-4.1, p=0.0003; Y coordinate: T=-1.5, p=0.14; Z coordinate: T=1.9, p=0.07; Left X coordinate: T=0.56, p=0.6; Y coordinate: T=-3.4, p=0.003; Z coordinate: T=1.6, p=0.12).

**Figure 3.**
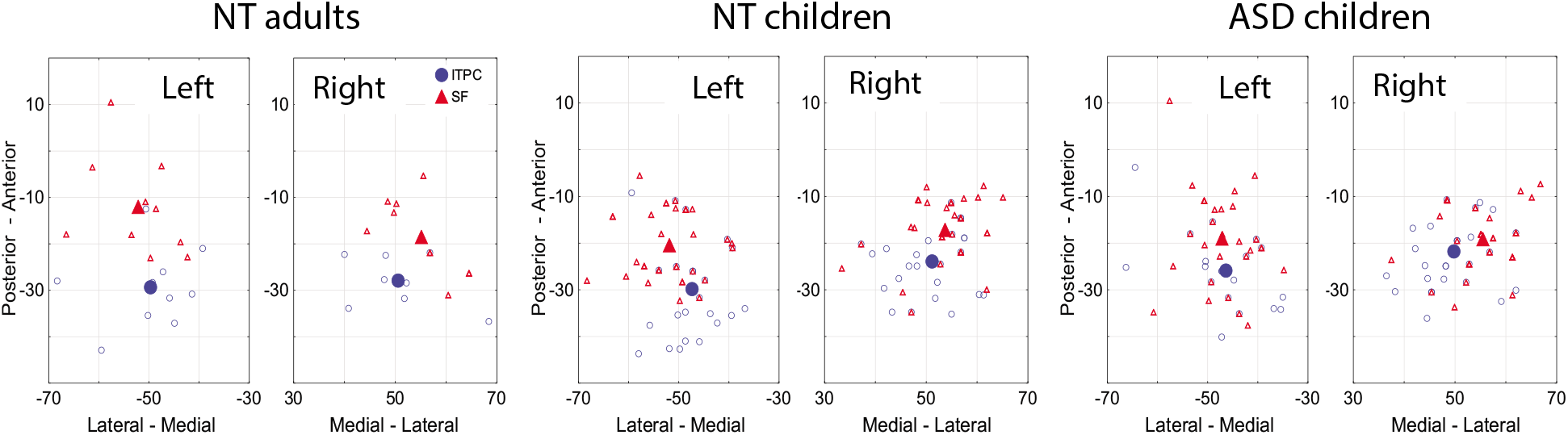
MNI coordinates of the 40Hz ASSR ITPC (blue shapes) and the SF (red shapes) maxima. Localization of the sources in the horizontal plane. Small open circles and small triangles correspond to individual coordinates; big filled shapes show the group means.

**Table 3.** Grand average MNI coordinates of the maximal ASSR and SF sources.

| Group<br>(N left / N right) | Left hemisphere Mean and (SD) |  |  | Right hemisphere Mean and (SD) |  |  |
| --- | --- | --- | --- | --- | --- | --- |
|  | X<br>(lateral-<br>medial) | Y<br>(posterior-<br>anterior) | Z<br>(superior-<br>inferior) | X<br>(medial-<br>lateral) | Y<br>(posterior-<br>anterior) | Z<br>(superior-<br>inferior) |
| **ASSR** |  |  |  |  |  |  |
| NT adults (10/10) | -49.7 (8.6) | -29.4 (8.5) | 9.9 (5.4) | 50.6 (8.1) | -28.0 (5.5) | 11.1 (4.3) |
| NT children (27/32) | -47.7 (6.0) | -28.4 (13.5) | 11.6 (7.5) | 51.2 (6.3) | -24.0 (6.3) | 10.4 (3.8) |
| ASD children (24/30) | -46.3 (7.6) | -25.9 (7.7) | 9.1 (4.7) | 49.7 (6.6) | -21.8 (10.4) | 9.6 (5.5) |
| *Keceli et al. 2015*<sup>#</sup> |  |  |  |  |  |  |
| Adults, N=11 |  |  |  |  |  |  |
| Estimated MNI | -48 | -22 | 11 | 52 | -17 | 13 |
| Original Talairach | -45 | -22 | 13 | 48.5 | 16.5 | 14 |
| **SF** |  |  |  |  |  |  |
| NT adults (10/10) | -52.1 (7.7) | -12.1 (10.7) | 1.2 (6.5) | 55.1 (6.8) | -18.5 (8.3) | 6.8 (6.2) |
| NT children (33/35) | -51.9 (6.7) | -20.5 (7.3) | 6.6 (3.7) | 53.7 (6.6) | -17.2 (6.6) | 7.4 (3.4) |
| ASD children (29/35) | **-47.1 (6.0)** | -19.0 (10.4) | 5.7 (7.4) | 55.5 (6.2) | -19.1 (6.5) | 7.5 (4.3) |
| *Keceli et al. 2015*<sup>#</sup> |  |  |  |  |  |  |
| Adults, N=12 |  |  |  |  |  |  |
| Estimated MNI | -51 | -17 | 7 | 52 | -13 | 6 |
| Original Talairach | -48 | -18 | 8.5 | 49 | -14 | 9 |

<sup>#</sup> The provided coordinates are approximated from the figure 3 in *Keceli et al. 2015*, where the authors localized ASSR and SF responses induced by the same periodic stimuli [20].
\* Significant difference from the NT children group: $p=0.005$ .

For the 40Hz ITPC maxima in the right or left hemispheres no significant differences between children with and without ASD were found in either X, Y or Z coordinates (T-test, all p’s>0.05). The SF maximum in the left hemisphere was located significantly more medial in children with ASD than that in NT children, although the difference was relatively small (X=coordinate in NT: −51.9, ASD: −47.1, F_(1,61)_=8.5, Cohen’s *d* = 0.74, p=0.005, uncorrected for multiple comparisons). The multivariate Hotelling T^2^ test confirmed presence of significant ASD vs NT differences in the source localization (F_(3,59)_=3.1, p=0.03; partial eta-squared = 0.14). No group differences were found for the SF coordinates in the right hemisphere.

To investigate whether the SF onset interval (150-250 ms), that was visible only in children, does represent the evolving SF, we compared cortical locations of the SF maxima in this interval with those in the 300–500 ms interval, where the sustained field was observed in both children and adults. No significant time-related differences in X, Y or Z SF coordinates were found in the NT, ASD or the combined sample of children in either hemisphere (paired T-test, p’s>0.08). This means that the onset interval of the sustained component in children originates from the same region of the auditory cortex as the rest of the SF.

### Comparison of the 40 Hz ASSR in NT children and children with ASD

To analyze the ASSR, we computed for each participant an average of 40 Hz ITCP values across 10 ‘maximally induced’ cortical vertexes, separately in the right and left hemispheres (see Methods for details). Grand averaged ASSR time-courses as well as the ITPC time-frequency plots for the left and the right auditory cortex are presented in Figure 4.

**Figure 4.**
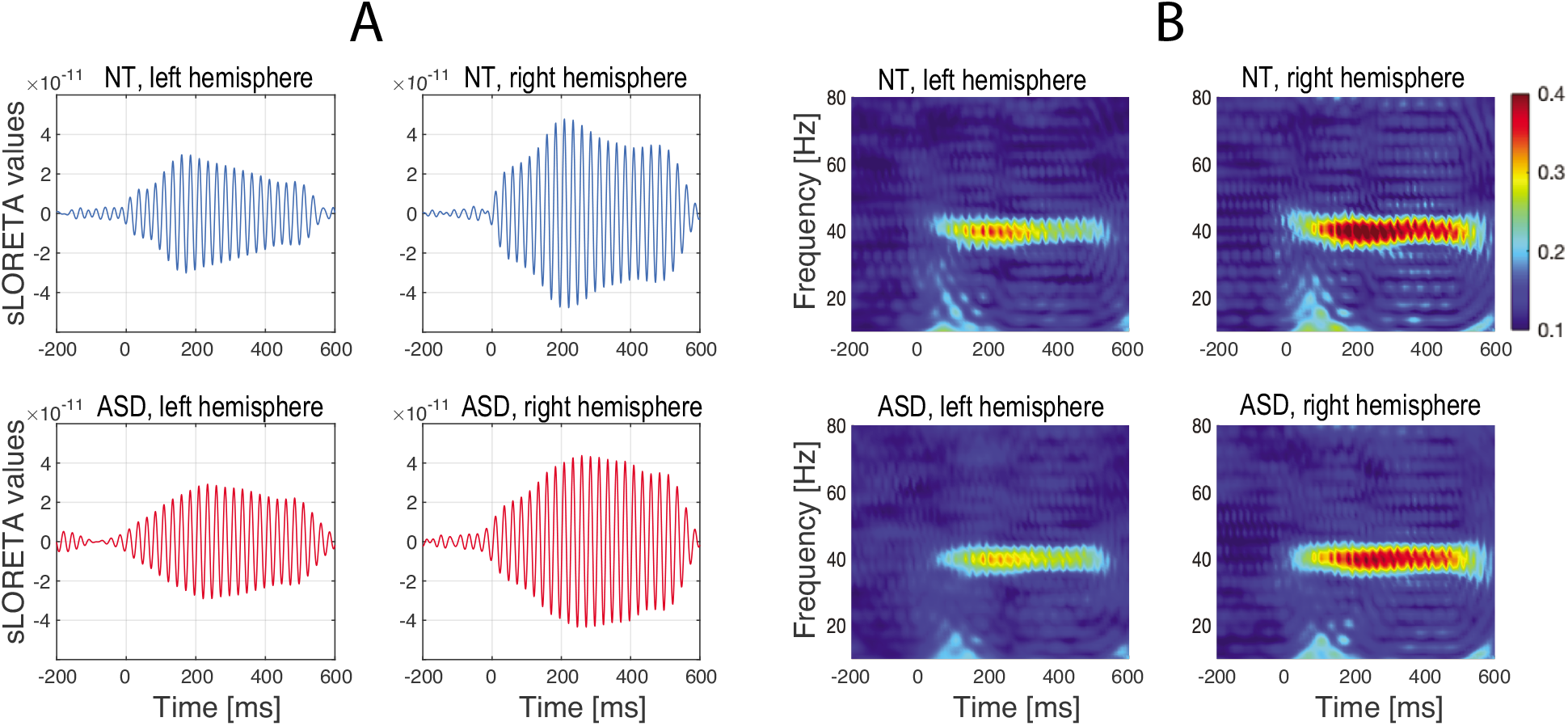
Grand average timecourses of the 40 Hz ASSR in the right and left response maxima in NT children and in children with ASD. Only responses to contralateral ear stimulation are shown. (A) Grand average ASSR filtered between 35-45 Hz. (B) Inter-Trial Phase Coherence (ITPC). No difference in the 40 Hz ASSR/ITPC between ASD and NT groups was found.

The contralateral 40 Hz ITPC values were approximately normally distributed in NT children (Shapiro-Wilk test, p’s>0.1) but their distributions significantly deviated from normality in children with ASD (Left hemisphere: Shapiro-Wilk W=0.83, p=0.00007; Right hemisphere: W=0.91, p=0.007). To normalize the distributions, the data were log10-transformed and the rmANOVA with factors Group, Hemisphere and Stimulation contralaterality (ipsi-vs contra) has been performed. The rmANOVA revealed highly significant effect of Hemisphere (F_(1, 68)_=41.7, partial eta-squared = 0.38, p<10e-7) and Stimulation contralaterality (F_(1, 68)_=54.3, partial eta-squared = 0.44, p<10e-9), but no effect of Group or its interactions (all p’s>0.05). In both groups of children, the 40 Hz ITPC was higher in the right hemisphere than in the left one and in response to the contra-relative to ipsilateral stimulation.

Figure 5 shows individual 40 Hz ITPC values in children with and without ASD as a function of age. The 40 Hz ITPC increased with age in both groups and in both hemispheres, contralateral to the stimulated ear. However, even at younger age (<9 years) the majority of children had 40 Hz ITCP values above their baseline level. This suggests that the majority of children had reliable ASSRs assessed by 40 Hz ITPC. Inspection of the residuals after subtraction of variability explained by age has shown that some children with ASD had very high for their age 40 Hz ITPC values (*Supplementary Figure S1, A and B*).

**Figure 5.**
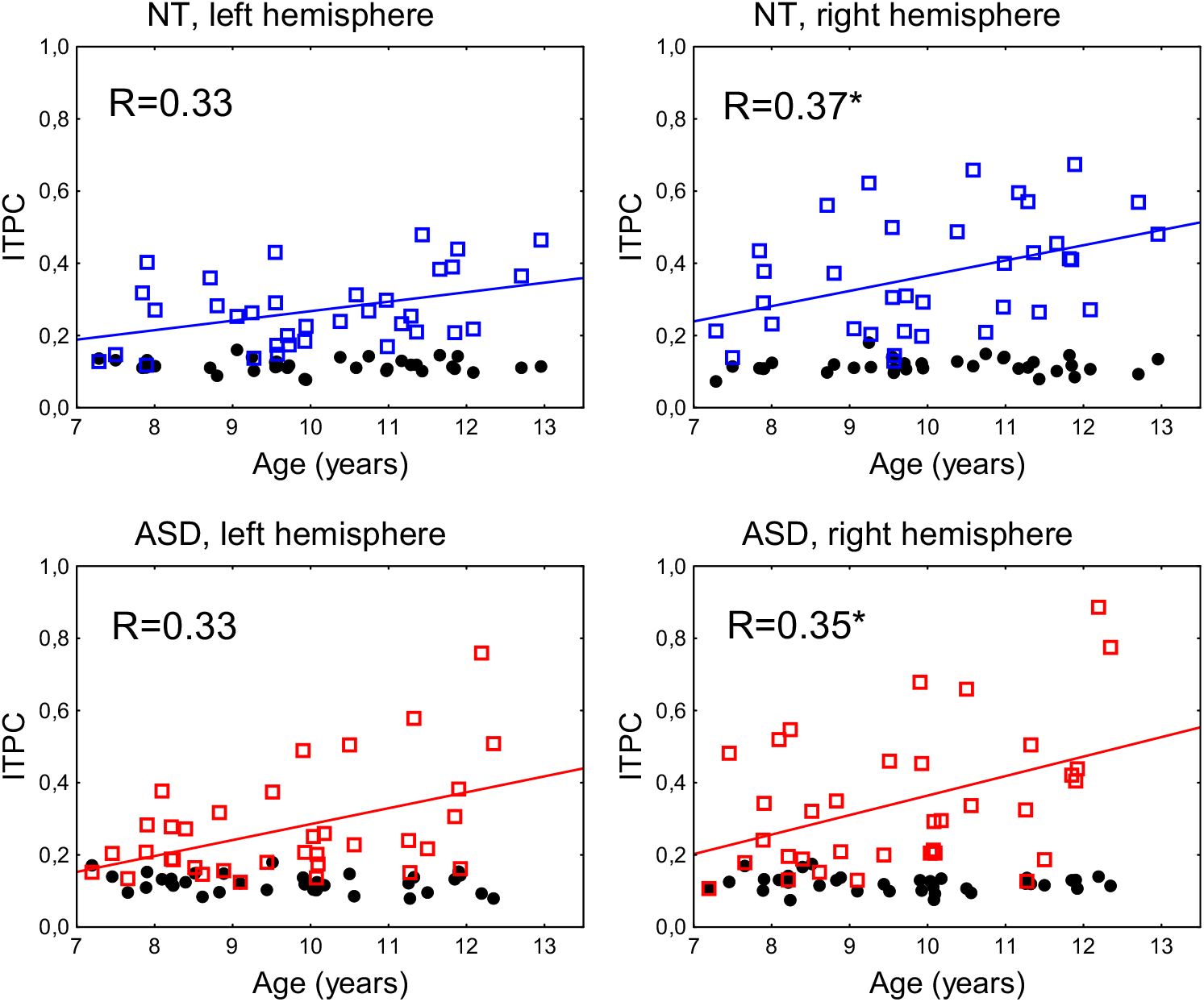
Associations between age and 40 Hz ASSR in NT children (upper panels) and children with ASD (lower panels). The individual stimulus-locked and baseline 40Hz ITCP values are shown in colored squares and black circles respectively. *R’s* are the Spearman correlation coefficients. Asterisk denotes significant correlation: p<0.05

To investigate whether the 40 Hz ITPC in children with ASD correlates with their intelligence level and severity of autism, we calculated Spearman correlations. Children with more severe autism had *higher* 40 Hz ITCP values in the right hemisphere (Table 4). Calculation of partial correlations for normalized (log-transformed) data, while controlling for age, increased reliability of this result (Age: partial *R*=0.46, p=0.008; Autism Score: partial *R*=0.45, p=0.009)

**Table 4.** Spearman correlation between the 40 Hz ASSR ITPC and psychometric variables in children with ASD.

|  | MPI IQ (N=32) | Autism Score (N=34) |
| --- | --- | --- |
| Left hemisphere | -0.11, $p=0.5$ | 0.25, $p=0.15$ |
| Right hemisphere | -0.14, $p=0.44$ | **0.40, $p=0.018^*$** |

\* Uncorrected for multiple comparisons

### 3.5. Comparison of the SF in children with and without ASD

As in case of the ASSR, between-group comparisons for the SF timecourses were performed in the source space (see Methods for details). Figure 6 shows the grand average high-passed SF source waveforms in the left and the right hemispheres for the contra- and ipsilateral ear stimulation in the ASD and the NT groups. Visually detectable SF source waveforms were present in the right hemisphere in all participants and in the left hemisphere in the majority of children from both samples, with the exception of the two NT and the six ASD children.

**Figure 6.**
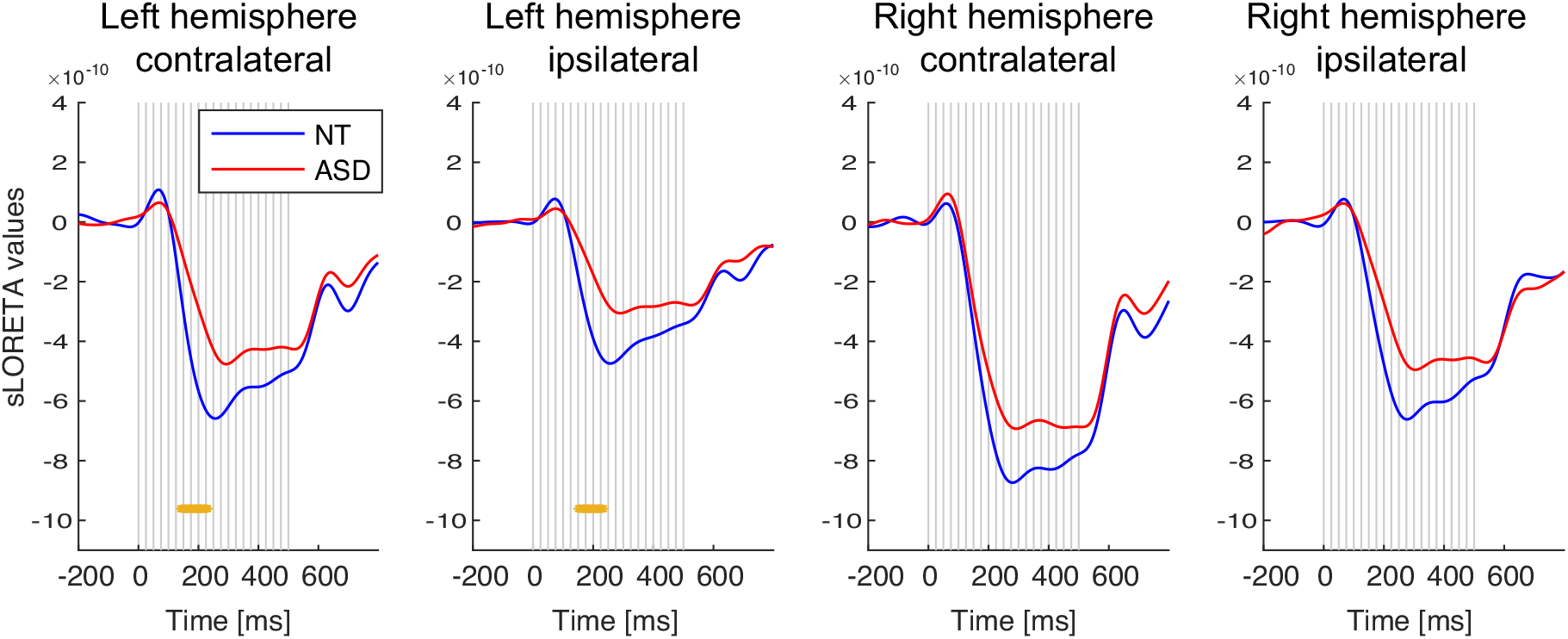
Comparison of the SF responses in the left and right cortical maxima in children with and without ASD. The signal was law-pass filtered at 9 Hz. Vertical gray lines mark clicks onsets. The yellow lines under the curves denote significant between-group differences on the point-by-point basis (Wilcoxon rank sum test, p<0.001, uncorrected for multiple comparisons).

First, we analyzed the effect of age on the SF maximal amplitude using Pearson correlations. None of the correlations was significant in the NT, ASD or in the combined sample (all p’s >0.15). This means that the maximal SF amplitude in boys does not change between 7 and 13 years of age.

Then we examined the effects of hemisphere and contralaterality of the stimulation on the SF maximal amplitude in the NT and ASD groups. To this end, we used rmANOVA with factors Group (ASD / NT), Hemisphere (Left / Right), Ear of Stimulation (contralateral / ipsilateral), and the maximal amplitude of the SF source current in the 150 - 500 ms stimulation interval as a dependent variable. There were strong effects of Hemisphere (F(1, 68)=50.5, partial eta-squared =0.43, p<,00001) and Ear of Stimulation (F(1, 68)=128.4, partial eta-squared =0.65, p<0.00001). The SF maximal amplitude was higher in response to the contra-than ipsilateral stimulation, and it was generally higher in the right hemisphere (Figure 6). None of interaction effects of the repeated-measure factors with Group were significant (all p’s>0.5), suggesting that the auditory SF response in both groups was characterized by contralaterality and the right hemisphere dominance. The SF maximal amplitude was higher in the NT than in the ASD group (main effect of Group: F(1, 68)=4.6, partial eta-squared =0.06, p=0.035). This means that the SF in children with ASD was reduced compared to NT controls in both hemispheres and in response to both contra- and ipsilateral ear stimulation. Analysis of the point-by-point group differences in the SF timecourses (Wilcoxon rank sum test, significance level was set to p<0.001) has shown that the SF in children with ASD was particularly strongly attenuated in the left hemisphere approximately between 150 ms and 220 ms after the click train onset, suggesting abnormally slow rise of the SF source strength in children with ASD (Figure 6).

For the further analysis of between-group difference in the SF source timecourse, we focused on the contralateral responses that were greater and more reliable than the ipsilateral ones (Figure 6).

To test for the group differences in the SF timecourses, we divided the SF time window (150 ms – 550 ms) into four successive time intervals in respect to the click train onset (151-250 ms; 251-350 ms, 351-450 ms and 451-550 ms), with the first (151-250 ms) interval corresponding to the raising part of the SF. We then performed rmANOVA with factors Time, Hemisphere and Group and the SF source current amplitudes in the 100 ms time intervals as a dependent variable (Table 5, Figure 7).

**Figure 7.**
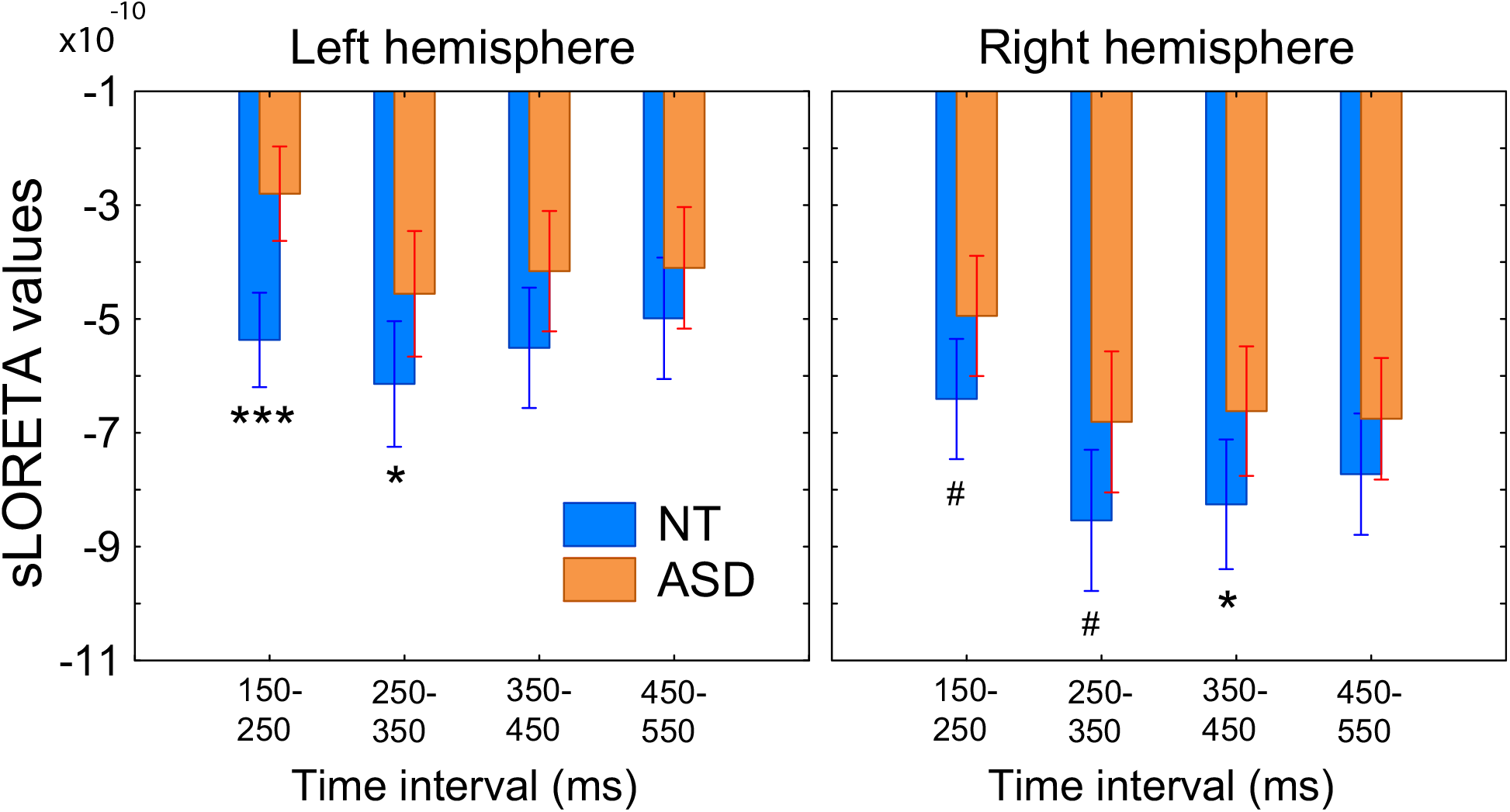
Group differences in the SF source amplitude in the consecutive temporal intervals. The SF responses in the left and the right hemisphere are evoked by stimulation of the contralateral ear. Asterisks indicate between-group differences. *p< 0.05, ***p <0.001, #p <0.1.

**Table 5.** Group differences in the SF source timecourses: rmANOVA results.

|  | *F* | *p* | *G-G*<br>*epsilon* | *partial*<br>*eta-squared* |
| --- | --- | --- | --- | --- |
| *Group* | **5.9** | **0.018** |  | 0.08 |
| *Time* | **32.4** | **<1e-6** | 0.57 | 0.32 |
| *Time x GR* | **3.32** | **0.04** | 0.57 | 0.05 |
| *Hemisphere* | **43.4** | **<1e-6** |  | 0.39 |
| *Hem x GR* | 0.04 | 0.84 |  | 0.00 |
| *Time x Hem* | **9.2** | **0.00001** | 0.71 | 0.12 |
| *Time x Hem x GR* | **3.78** | **0.024** | 0.71 | 0.05 |

Children with ASD had generally lower SF source amplitude than NT children, especially during the rising part of the SF timecourse – from 150 ms 250 ms after stimulation onset (*Time x GR* interaction: p<0.05; Figure 7). The significant *Time x Hem x GR* interaction (p=0.024) was due to a greater and more reliable reduction of the ‘early’ SF segment in the left hemisphere than in the right one in children with ASD (ASD vs NT left hemisphere: p <0.001; right hemisphere: p<0.1). This result is consistent with that presented in Figure 6.

To check if the group differences could be explained by ‘developmental delay’ we calculated Pearson correlations between age and the averaged SF source amplitudes in the four time intervals in the left and right auditory cortices. No correlations with age were found for the SF source amplitudes in either the NT or the ASD group (all uncorrected p’s >0.19).

Considering that there were small, but significant group differences in the SF localization in the left hemisphere (Table 3), we wanted to ensure that the between-group differences in the SF source amplitude were not driven by the choice of the SF vertices for the group analysis, which was done by averaging across the NT and ASD data (see Methods for details). For this purpose, we repeated the rmANOVA analysis for the SF source amplitude calculated in the individually chosen 30-vertices with maximal SF amplitudes. The results remained principally the same (GR: F_(1,68)_=3.7, partial eta-squared=0.05, p=0.057; Time x GR: F_(1,68)_=8.9, partial eta-squared=0.05, p=0.023; Time x Hem x GR: F_(3,204)_= 5.6, partial eta-squared=0.08, p=0.003, epsilon=0.76)

To summarize, in both the NT and the ASD children, the SF source strength was higher contralaterally to the stimulated ear and clearly dominated in the right hemisphere. In the ASD group, the SF was moderately reduced in both hemispheres and delayed in the left one, irrespectively of the stimulated ear. None of the SF parameters changed with age between 7 and 12 years in either the NT or the ASD group.

We further performed correlation analysis to investigate whether the two principle findings – the bilateral reduction of the SF maximal amplitude and the left-hemispheric SF reduction in its 150-250 ms onset interval (‘SF delay’) in children with ASD – are associated with psychometric variables (Table 6).

**Table 6.** Partial correlations between the SF source current* and psychometric variables in children with ASD (N=31).

| SF amplitude | MPI IQ: r, p* | Autism Score |
| --- | --- | --- |
| $SF_{max}$, Left hemisphere | -0.35, $p=0.06$ | -0.22, $p=0.25$ |
| $SF_{max}$, Right hemisphere | **-0.43, $p=0.017$** | **-0.64, $p=0.0001$** |
| $SF_{150-250}$, Left hemisphere | -0.06, $p=0.74$ | -0.04, $p=0.85$ |
| $SF_{150-250}$, Right hemisphere | -0.14, $p=0.47$ | **-0.41, $p=0.026$** |

\* Note, that the SF current is negative, and negative correlation reflects a direct link between the SF strength and a respective psychometric variable
\*\* Uncorrected for multiple comparisons

Since children with lower MPI IQ scores had higher severity of autism (Pearson *R*=-0.36, p= 0.049), to estimate independent associations of these psychometric variables with the SF amplitude, we calculated partial correlations. This analysis has been performed in 31 children with ASD in whom both MPI IQ and Autism Scores were available (Table 6). Although the maximal SF source amplitude (SF_max_) was decreased (less negative) in children with ASD at the group level, its greater strength in the right hemisphere correlated with *greater* severity of autism traits. The similar correlation was observed for the SF onset interval (SF_150-250_) in the right hemisphere. When corrected for autism severity, the SF_max_ (but not SF_150-250_) in both hemispheres tended to be higher in the ASD participants with higher IQ. Thus, the results of the correlation analysis mainly indicate that higher SF amplitude in the right hemisphere in children with ASD is associated with greater severity of their autism symptoms.

## Discussion

We investigated putative hemispheric differences in low-level cortical processing of periodic spectrally-complex sound in boys with ASD and age-matched control boys. To this end, we used 40 Hz click trains presented separately for the left and right ears to probe contralateral auditory responses. In children, similarly to adults, temporally regular clicks evoked two types of sustained neural responses with different sources alongside the Heschl’s gyrus: the ASSR and SF. Regardless of age, the SF sources in both hemispheres were positioned anterolateral to those of the ASSR. These results agree with the previous report that has shown similar relative positions of neural generators of these components in adults [20]. While the ASSR in children with ASD did not distinguish them from their typically developing peers, their SF was reduced in both hemispheres, as well as markedly delayed and spatially displaced in the left hemisphere. In this section we discuss the implications of these results in the context of previous neuroimaging and neuronal studies of temporal regularity processing in the auditory cortex.

### ASSR in NT children and children with ASD

In accordance with the previous findings in adults [21, 57], the monaural stimulation in children elicited the ASSR with significant contralateral predominance and generally higher magnitude in the right hemisphere (Figure 4). In both children and adults, the ASSR was localized in or in vicinity of the primary auditory cortex (area A1) (Figure 3). Despite differences in source localization techniques, the MNI coordinates of ASSR maxima in our study were similar to those previously reported by studies performed in adults using single dipoles anatomically constrained to the left and right Heschl’s gyri [20, 57].

The phase consistency of the ASSR – the most individually reliable feature of this type of response to periodic clicks [53] – increased between 7 and 13 years of age (Figure 5), which is in line with the available evidence of marked enhancement in the 40-Hz ASSR between late childhood (8–10 years) and early adolescence [29, 58–60]. This age-related increase can be explained by developmental changes in fast-spiking parvalbumin-sensitive neurons involved in generation of sensory gamma oscillations and their entrainment by periodic auditory stimuli [58].

In the majority of children in our study the 40 Hz ITPC values were clearly above the baseline (Figure 5). This result contrasts with that of Edgar et al [29] who found that the majority of 48 NT children aged 7-14 years lacked a discernible 40 Hz ITPC responses. The higher proportion of children with reliable ASSR in our study can be explained by methodological differences, such as the use of clicks instead of tones, monaural instead of binaural stimulation, source localization methods, etc. Whatever the reason, our paradigm allowed us to detect ASSRs in most children. This is important, because Edgar et al suggested that inaccurate ASSR measurement during childhood might preclude the observation of atypical ASSR in the pediatric populations with neuropsychiatric disorders, such as ASD.

Despite the improved SNR, similarly to Edgar et al [29] and Ono et al [22], we found no significant difference in ASSR between the ASD and NT groups of children (Figure 4). Also in accordance with the Edgar et al study, children with ASD demonstrated a normal developmental increase in ASSR in both hemispheres (Figure 5), which strengthens the conclusion that the ASSR matures normally between 7 and 14 years of age in children with ASD, at least at the group level. Our findings and those of Edgar et al differ from the results of Wilson et al [25], who observed a reduced left-hemispheric 40 Hz ASSR to monaural clicks in 10 children with ASD as compared to 10 NT children. We should note, however, that the small sample size in this frequently cited research reduces the likelihood that a statistically significant result reflects the true effect [61]. Overall, our results demonstrate fairly typical development of the ASSR in the primary auditory cortex in children with ASD, at least until puberty. In this context, abnormal reduction of the ASSR found in several studies in adolescents and adults with ASD [23, 24, 28, 62] may indicate that the maturation of the primary auditory cortex in ASD diverges from the normal trajectory during or after puberty.

Distribution of the 40-Hz ITCP values in children with ASD significantly differed from normal, unlike that in the NT children. It is noteworthy that some children with ASD had ITPC values that were very high for their age (*Supplementary Figure S1, A and B*). Moreover, the magnitude of the 40-Hz ITCP in the contralateral right hemisphere was *higher* in those children with ASD who had more severe behavioral symptoms of autism (Table 4). Although, given the results of previous ASD studies, this was an unexpected finding, it is consistent with a direct correlation between the magnitude of 40 Hz ASSR and severity of positive symptoms in patients with schizophrenia [63]. It is likely that one of the factors that increase ASSR is the high excitability of the auditory cortex. It has been suggested that an increased propensity for high-frequency synchronization in the primary auditory cortex indicates its hyper-excitable state [64]. This hypothesis is compatible with our findings as well as with the animal data on augmented ASSR under high arousal states [64].

### Sustained Field in NT children and adults and its abnormalities in children with ASD

We found that in children, similarly to adults [18], the sustained neural response to temporally regular 40 Hz clicks consists of two superimposed neural signals: the ASSR and slowly developing negative DC shift, referred to as the Sustained Field. The current of the SF source has the same direction in both age groups (Figure 2), which speaks in favor of functional homology of the SF in children and adults. It has been suggested that in adults the SF onset is hidden because it overlaps with the sequence of obligatory transient responses to the stimulation onset - P50m, N100m and P200m [18]. The same is probably true in children. The seemingly earlier onset of SF in children than in adults might be explained by the absence of obligatory components with longer latencies (N100m and P200m) in the immature auditory evoked field (Figure 2, see also [36–38]). Apart from difference in the onset time, the properties of SF in children are similar to those in adults.

*Firstly*, distributed localization modeling shows that in both children and adults, SF maxima are located in or in vicinity of the core of the auditory cortex, contralateral to the stimulated ear and anterolateral to the ASSR maxima (Figure 3; Table 3). The results of MEG inverse solutions depend on many factors, including the forward models used [65]. Therefore, the exact MEG-derived coordinates of the SF and ASSR sources should be treated with caution. Nevertheless, since the *same relative position* of the ASSR and SF sources has been reported by Keceli et al, who used localization technique different from ours - single dipole source modeling [20], we are confident in the validity of these results. Besides, the group average coordinates of the SF maxima in our participants are remarkably similar to the coordinates of the ‘anterior SF source’ derived from a four-dipole model in the study of Gutschalk et al. (see table 1 in [18]). It is therefore likely that, similarly to adult, the SF in children is generated in the pitch processing center [19, 33, 57] located immediately anterolateral to the primary auditory cortex in the lateral Heschl’s gyrus [12, 16].

*Secondly*, the dependence of the SF strength on the stimulated ear is identical in children and adults – in both groups the SF has greater amplitude in the hemisphere contralateral to the stimulated ear, and, in general, in the right hemisphere (Figure 2). The same hemispheric asymmetry was previously described for adults [21].

Unlike the ASSR, the SF in children with ASD was clearly abnormal. This dissociation between the two complementary neural responses to periodical clicks provides a clue to the lowest level of the cortical hierarchy at which the periodicity processing may be impaired in children with ASD.

The temporal regularity at a frequency of 40 Hz is explicitly represented in monkeys’ area A1 by neuronal firing patterns synchronized with each individual click [66]. The human ASSR, most probably, reflects this ‘stimulus-synchronized code’ implemented in area A1 [67]. However, the ASSR is thought to reflect neural coding of relatively slowly repeating acoustic events (up to 50 Hz) [67]. The tonotopically-organized A1 neurons, although involved in coding of pure tones over a wide range of frequencies, are limited in their capacity to synchronize with the periodic modulations in spectrally complex sounds at rates faster than about 100 Hz [66].

The click repetition rate of 40 Hz, which is above the lower frequency limit for perceiving click train as a continuous sound (approximately 30 Hz; [8]), also activates the second neural code for temporal regularity – the monotonous rate-coding [9, 11, 66]. Perception of pitch in harmonic continuous sounds corresponds to the lowest rate at which the periodic waveform repeats itself, and the inverse of this period is called the fundamental frequency (*f_0_*) [44]. It has been shown that in monkeys the increasing *f_0_* of the click train is reflected in a monotonous increase in the firing rate of neurons located in the cortical field rostral to the A1, which is homologous to the human pitch-processing center in the auditory core area [9]. Given that the neuronal firing rate in cortical evoked responses is associated with surface-negative DC shift in non-invasive EEG/MEG (see [68] for review), it is conceivable that the SF reflects activity of the rate-coding neurons in the pitch-processing center [57]. Indeed, similarly to neuronal firing rate in monkeys, the anterior SF power in humans is directly proportional to the *f_0_* of a click train and is strongly sensitive to violations of temporal periodicity, both in click trains and in human vowels [33]. Gustchalk et al argued that the anterolateral portion of the Heschl’s gyrus – the area that generates the anterior SF – serves to integrate pitch information across different frequency channels and/or calculates the specific pitch value in spectrally complex sounds.

Based on these considerations, the coexistence of normal ASSR (Figures 6, 7) with reduced and delayed SF (Figures 4, 5) in our participants with ASD is consistent with the emerging picture of dual temporal encoding of 40 Hz fundamental frequency in the human auditory core [9, 67]. While the ‘stimulus-synchronized code’ employed by the primary auditory cortex seems to be relatively spared in children with ASD, the ‘rate-coding’ implemented in the anterolateral region of the Heschl’s gyrus might be disrupted.

The undisturbed functioning of the primary auditory cortex in children with ASD is well in line with numerous electrophysiological evidence on their typical, or even enhanced processing of frequency/pitch of pure tones [69–71] for review) that mainly relies upon A1 neural circuitry. On the other hand, the SF abnormalities suggestive of abnormal functioning of the ‘pitch processing center’ in our participants with ASD, generally agree with the conclusion derived from the comprehensive meta-analysis of mismatch-negativity (MMN) studies in autism [71]. The results of this meta-analysis point to a weak neural encoding of *spectrally and/or temporally complex non-speech stimuli* in the prepubertal children with ASD. However, the previous MMN research clarified neither the cortical level at which this deficit occurs nor its hemispheric lateralization in children with ASD.

In this respect, the most important and intriguing finding of the current study is the left-hemispheric preponderance of the SF abnormalities in children with ASD. The SF was markedly delayed in the left hemisphere in children with ASD, while its maximal strength was slightly, but significantly, reduced in both hemispheres in children with ASD (Figures 6, 7). Moreover, the source of SF maximum in the left hemisphere in children with ASD was located more medially than that in the control group.

There are several potential factors that could obscure the measurements of auditory evoked fields in children with ASD, but they can hardly explain their SF abnormalities found in our study. Poor quality of MEG recording in children with ASD (see e.g. [71] for discussion) is unlikely to explain the group differences. Indeed, the ASSR phase consistency did not differ between the ASD and NT groups (Figure 4), pointing to comparable SNR. Hampered involuntary deployment of attention to salient auditory stimulation, such as e.g. phonemes in children with ASD (see e.g. [72, 73] for discussion), can also be refuted due to a neutral, purely sensory nature of the stimulation in our study. Besides, differences in attentiveness to the flow of auditory stimulation would equally affect AEFs in both hemispheres and cannot explain the observed left-hemispheric SF deficit in children with ASD. Differences in automatic orienting to an abrupt onset of click stimulation in ASD [36] affect the auditory responses predominantly in the right hemisphere [74–76] and could not elicit selective left hemispheric delay of SF in our ASD participants. The delayed maturation of the N100m in the clinical group could affect the activation strength around 150-200 ms after the click train onset, because the N100m has the same direction of current as the SF and, if present, partially overlaps with its early segment. However, the N100m matures earlier in the right than in the left hemisphere [36, 37] and its delayed maturation would rather affect the right-hemispheric SF in children with ASD. The lack of significant correlations between amplitude of the SF in its ‘early’ segment (150 to 250 ms) and age makes this explanation even more unlikely. To sum up, the abnormally slowly evolving SF in the left hemisphere in children with ASD appears to be a true phenomenon and is not a result of the ‘developmental delay’ or a methodological artifact.

Remarkably, we observed a small but significant group difference in the source location of the in the left but not in the right hemisphere. Previous studies in people with ASD reported morphological abnormalities in language-related areas of the left temporal lobe [77–79], but also in Heschl’s gyri [80]. We did not perform structural brain analysis, and it is therefore unclear if the left-hemispheric difference in SF localization reflects primarily functional or morphological differences between the groups. Nevertheless, this finding is generally in line with our hypothesis regarding the dysfunction of the left-hemispheric pitch-processing center in ASD.

The functional relevance of the left-hemisphere SF abnormality in children with ASD can be explained in light of the different functional roles of the left and right anterolateral regions of the auditory core in processing the fundamental frequency/pitch of spectrally complex harmonic sounds. Periodicity in complex harmonic sounds is analyzed in the auditory system through two mechanisms that play different but complementary roles (for review see [9]). One of them performs rapid extraction of the *f_0_*, i.e. the repetition rate of the temporal envelope, while another relies upon spectral calculation of the relationships between harmonics (*f_sp_*). The latter mode is more precise and universal as it works even if the *f_0_* is not physically present in the stimulus. The 40 Hz click trains used in our experiment were characterized by the presence of both *f_0_* and its higher frequency harmonics (Figure 1), which implies the involvement of both mechanisms in the processing of this signal. Although it is generally accepted that pitch processing in music and speech is lateralized to the right hemisphere (e.g. [81]), a number of neuroimaging and lesion studies suggest a stronger sensitivity of the right hemisphere to spectral properties of periodic sounds (for review see [82]), and a relative left-hemispheric specialization for rapid decoding of its fundamental frequency [83, 84]. In particular, a larger grey matter volume of the left anterolateral Heschl’s gyrus predisposes people to hear the *f_0_* in the ambiguous sound, while a greater volume of this region in the right hemisphere inclines them to rely on the *f_sp_* when processing pitch [16].

There is strong evidence that such low-level left-hemispheric specialization for processing of the temporal envelope is directly associated with the capacity to incorporate pitch information into perception of speech [17]. Wong et al reported that people who were more successful at learning how to use pitch difference to discriminate unknown pseudowords and associate them with visual images, also had greater grey matter volume in the left Heschl’s gyrus (but not in the right Heschl’s gyrus).

Given the above, we suggest that selective slowing of neural activation in the left anterolateral Heschl’s gyrus in children with ASD may hinder their linguistic pitch processing, even if their perception of musical pitch is preserved. Although speculative, this explanation is consistent with the paradoxical dissociation between undisturbed or even enhanced ability to perceive melodic contour (pitch changes) in non-speech stimuli in people with ASD [85, 86], and their difficulties with decoding and producing pitch variations (e.g. intonation) in speech [87, 88]. Interestingly, this dissociation is present even in tone language speakers with autism, suggesting that the tone language experience does not compensate for their speech intonation perception deficit [89]. The difficulties with linguistic pitch processing and production in people with ASD are relatively independent from their intelligence, severity of communication disturbances and even from the other language skills [87, 90]. This may explain absence of correlation between the ‘early’ SF and psychometric variables in our study (Table 6).

At this point, it is unclear whether the lateralized SF abnormalities observed in children with ASD in our study are specific for processing of the temporal regularities in sound or reflect generally disturbed left hemisphere specialization across multiple cortical areas that has been found in ASD individuals in structural MRI studies [91, 92].

In contrast to the left-hemispheric delay in the SF amplitude growth, its maximal amplitude was slightly, but significantly, reduced in both hemispheres in children with ASD (Figure 7). Unexpectedly, in the right hemisphere both greater SF strength and higher 40 Hz ITPC were associated with *greater* severity of behavioral symptoms of autism (Table 4, and 6). Both these correlations can be explained by inter-individual variations in a certain factor, such as increased neural excitability of the right auditory cortex, that non-specifically affects the auditory evoked responses in some children with ASD.

To summarize, the monaural presentation of the periodic 40 Hz click trains evokes typical ASSR, but abnormal SF response in children with ASD. The SF, which was generated in the Heschl’s gyrus, anterolateral to the ASSR source, was bilaterally reduced and strongly delayed in the left hemisphere contralateral to the stimulated ear in children with ASD. Functionally, the anterolateral part of the Heschl’s gyrus plays a role in processing the pitch of harmonic complex sounds, including speech. The abnormally slow build-up of the SF response associated with pitch processing was present in children with ASD irrespective of their IQ and severity of autism symptoms. In this respect, this neuro-functional deficit resembles the well-known difficulties with perception and production of *f0* in speech intonation, which is also present in people with ASD regardless their IQ and autism severity. We assume that deficient low-level processing of fundamental frequency in the ‘pitch processing center’ of the left hemisphere may contribute to the abnormal perception and production of speech prosody in people with ASD.

## Limitations

Our study has several important limitations. Firstly, our findings are confined to 7-12 year-old children. Given the abundant evidence of dramatic developmental changes in the auditory cortex during puberty, future studies should clarify how both types of auditory neural responses to temporal regularity – the ASSR and SF – change during and after this age, and whether their development trajectories in people with ASD begin to deviate from the typical ones after puberty. Secondly, our stimulation protocol did not directly contrast regular interval noise and aperiodic noise stimuli (see e.g. [19]). Therefore, our statement about sensitivity of the child SF to stimulus periodicity is based on previous MEG studies in adults and needs to be tested experimentally in the future studies. Thirdly, since we used only 40 Hz click trains, we cannot completely rule out the possibility that the SF left-hemispheric abnormality in the ASD population is confined to this repetition rate, which is only slightly above the lower limit of pitch perception [8]. Fourthly, in the present study we did not test the ability of our participants to process vocal and non-vocal pitch. In future studies we plan to combine the SF measurements with assessment of prosody perception and production in children with ASD.

## Conclusions

Our study provides the first evidence of specific left-hemispheric abnormality in the sustained neuromagnetic response to 40 Hz click trains in children with ASD. Given that this neural response is associated with processing of sound envelope periodicities pertinent to pitch perception, we suggest that its left hemispheric deficit may be related to selective difficulties with pitch processing in speech context experienced by children with ASD. Our findings appear to be the first to show that impaired specialization of the left auditory cortex in children with ASD is not restricted to the high-level abstract representations of vocal speech, but occurs already at the low-level processing stage mainly concerned with extracting temporal regularity in a time varying acoustic signal.

## List of abbreviations

AEF: auditory evoked field
ASD: autism spectrum disorders
ASSR: auditory steady state response
ITPC: inter-trial phase coherence
MEG: magnetoencephalography
NT: neurotypical
SF: sustained field

## Funding

This work has been supported by the research grant from The Moscow State University of Psychology and Education (MSUPE).

## Authorship

### Author contributions

S.T.A. and O.E.V. designed research; G.D.E, O.T.S, O.T.M., P.A.O. performed research; O.E.V. and K.K.S. analyzed data; and S.T.A. and O.E.V. wrote the paper.

### Data availability

The datasets analysed in the current study is available from the corresponding author on reasonable request.

## Notes

### Competing Interest Statement

The authors have declared no competing interest.

