## Supplementary Figure S1 for "Left hemispheric deficit in the sustained neuromagnetic response to periodic click trains in children with ASD"

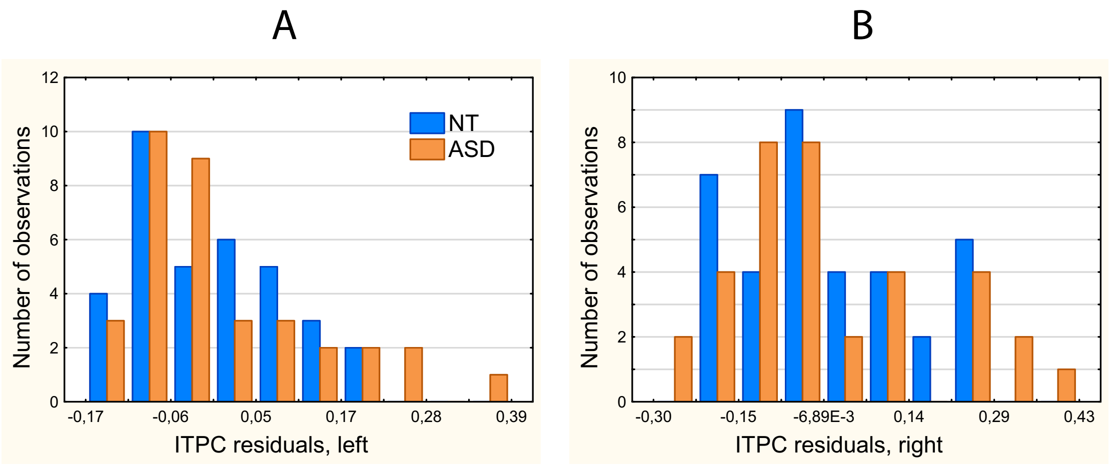


**Figure S1.** ITPC residuals in NT and ASD children after regressing-out age. (A) Left hemisphere. (B) Right hemisphere. Note presence of very high for their age ITPC values in some participants with ASD.
